# Unravelling the progression of the zebrafish primary body axis with reconstructed spatiotemporal transcriptomics

**DOI:** 10.1101/2024.07.01.601472

**Authors:** Yang Dong, Tao Cheng, Xiang Liu, Xin-Xin Fu, Yang Hu, Xian-Fa Yang, Ling-En Yang, Hao-Ran Li, Zhi-Wen Bian, Naihe Jing, Jie Liao, Xiaohui Fan, Peng-Fei Xu

**Author notes:** Corresponding authors: J.L., X.F. and P.X. Yang Dong, Tao Cheng and Xiang Liu contributed equally to this work and should be regarded as joint first authors.

## Abstract

Elucidating the spatiotemporal dynamics of gene expression is essential for understanding complex physiological and pathological processes. Traditional technologies like *in situ* hybridization (ISH) and immunostaining have been restricted to analyzing expression patterns of a limited number of genes. Spatial transcriptomics (ST) has emerged as a robust alternative, enabling the investigation of spatial patterns of thousands of genes simultaneously. However, current ST methods are hindered by low read depths and limited gene detection capabilities. Here, we introduce Palette, a pipeline that infers detailed spatial gene expression patterns from bulk RNA-seq data, utilizing existing ST data as only reference. This method identifies more precise expression patterns by smoothing, imputing and adjusting gene expressions. We applied Palette to construct the ***<u>D</u>anio <u>re</u>rio* <u>S</u>patio<u>T</u>emporal <u>E</u>xpression <u>P</u>rofiles (*Dre*STEP)** by integrating 53-slice serial bulk RNA-seq data from three developmental stages with existing ST references and 3D zebrafish embryo images. *Dre*STEP provides a comprehensive cartographic resource for examining gene expression and spatial cell-cell interactions within zebrafish embryos. Utilizing machine learning-based screening, we identified key morphogens and transcription factors (TFs) essential for anteroposterior (AP) axis development and characterized their dynamic distribution throughout embryogenesis. In addition, among these TFs, Hox family genes were found to be pivotal in AP axis refinement. Their expression was closely correlated with cellular AP identities, and *hoxb* genes may act as central regulators in this process.

## Introduction

Model organisms such as zebrafish have long been valuable tools for studying developmental biology and human diseases. Understanding the spatiotemporal patterns of gene expression in these models is crucial for gaining insights into the physiological and pathological mechanisms in normal development and related diseases. Thus, great efforts are ongoing to construct gene expression maps of these models with higher resolution, depth, and comprehensiveness.

Traditional technologies, such as ISH and immunostaining, have been widely used for investigating the spatiotemporal expression patterns of specific genes. However, these approaches are limited in their ability to simultaneously detect the expression of a large number of genes. In recent years, significant progress has been made in developing technologies for obtaining transcriptomics with spatial information. Techniques such as laser capture microdissection/microscopy (LCM) combined with bulk RNA-seq^1, 2^, Tomo-seq^3^, and Geographical positional sequencing (Geo-seq)^4^ have allowed the generation of spatially resolved transcriptomic data^5^. Additionally, methods like seqFISH^6^, MERFISH^7^, Slide-seq^8^, 10x Visium^9, 10^, and Stereo-seq^11–13^ have further improved the spatial resolution.

While these spatial transcriptomics (ST) techniques have advanced the spatial resolution of transcriptomic data, bulk RNA-seq remains the preferred choice for most studies due to limitations associated with ST techniques such as low read depth, suboptimal gene detection capability, and high cost^5, 14^. Consequently, tools have been developed to infer cell features or spatial gene expression from bulk RNA-seq data, including TIMER^15^, MuSiC^16^, DWLS^17^, and Bulk2Space^18^.

In this study, we introduce Palette, a pipeline designed to allocate gene expression from bulk RNA-seq data to spatial spots using ST data as the only reference. Palette has demonstrated its effectiveness in inferring spatial expression patterns in both *Drosophila* and zebrafish sections. We performed bulk RNA-seq on serial cryosections of zebrafish embryos along the left-right axis at three developmental stages. By applying Palette to the obtained data with the Stereo-seq data^12^ as references, we inferred the spatial gene expression patterns. We then projected the constructed 3D ST maps onto the zebrafish embryo images with 3D coordinates^19^ to correct the deformation during cryosectioning and construct a 3D spatial gene expression cartograph that more accurately reflects embryonic morphology. We named this cartograph *Dre*STEP, which enables the visualization of gene expression patterns in the context of the 3D morphology of the zebrafish embryos. Finally, leveraging the capabilities of *Dre*STEP, we characterized potential roles of morphogens and TFs in AP refinement during the progression of the primary body axis.

## Results

### Design concept of Palette

The overall working pipeline of Palette is depicted in Figure 1, illustrating the key steps involved in our approach. The pipeline firstly incorporates spatial clustering and deconvolution processes to account for differences in cluster abundances between bulk RNA-seq and ST data. Then, a variable factor is introduced to adjust expression differences between the two types of data. Subsequently, the pipeline estimates gene expression in each spot using a loop algorithm that takes into account regional gene expression, spot characteristics, and spot-spot distances. This iterative process allows for the inference of spatially resolved gene expression from bulk transcriptome data with relatively stable gene expression. The pipeline outlined in Figure 1 represents the sequential steps employed in Palette to accurately allocate gene expression to spatial spots using the information provided by the bulk RNA-seq data.

**Fig. 1.**
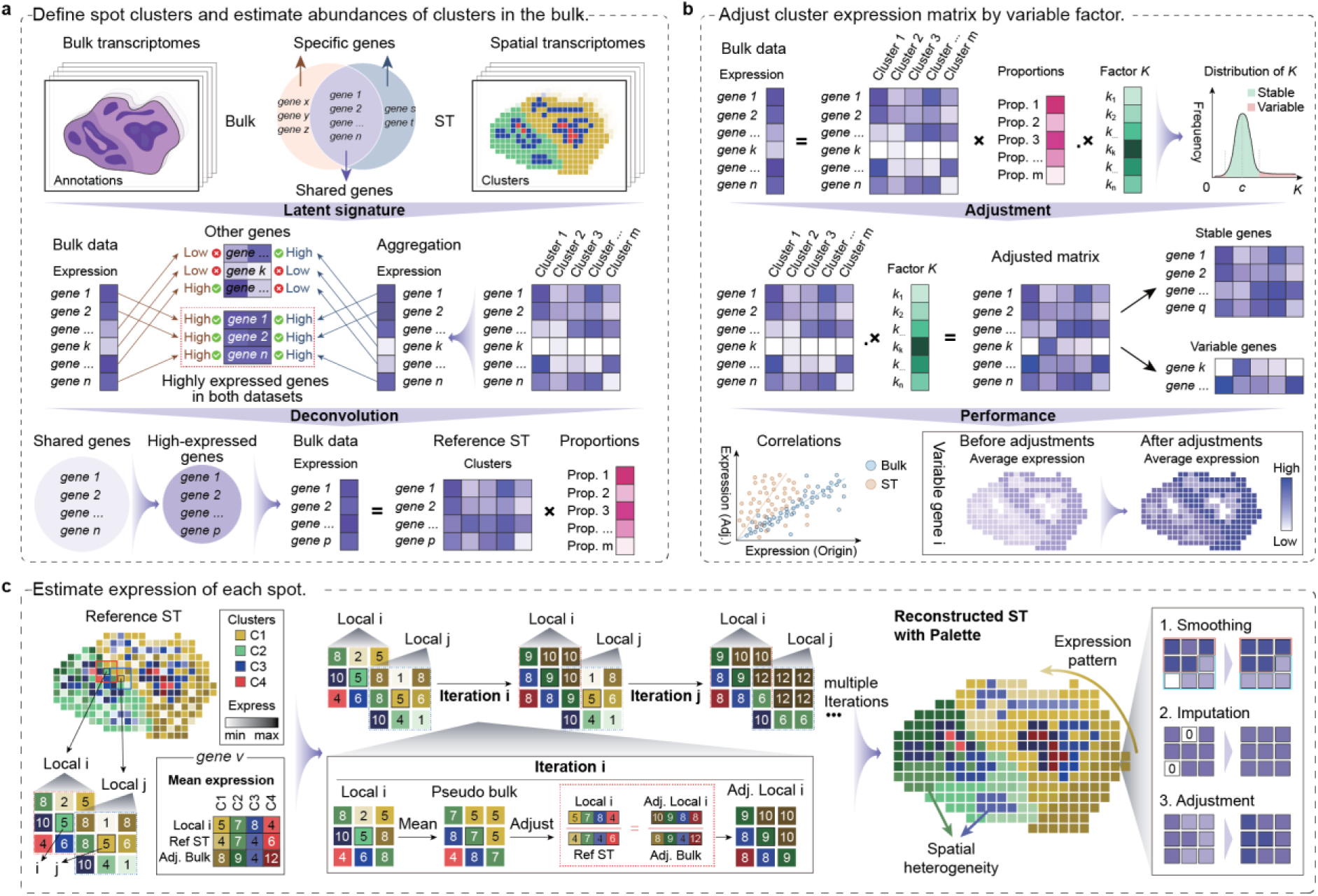
Working pipeline of Palette. **a,** Defining spot clusters in ST reference and estimating cluster abundances in bulk transcriptome data. Bulk transcriptome data and ST reference are taken as input. Highly expressed genes in both datasets are used for spatial clustering through BayesSpace^20^. The cluster expression matrix obtained from spatial clustering is then used as the reference for performing deconvolution on bulk transcriptome data, resulting in the estimated cluster abundances of bulk transcriptome data. **b,** Adjusting cluster expression matrix by employing the variable factor. The variable factor represents the expression differences between bulk RNA-seq data and ST reference. **c,** Estimating the expression in each spot through a loop algorithm. The expression of LST is adjusted and then evenly allocated to each spot of this cluster. After the looping steps, the average expression of each spot is taken as the estimated expression.

The detailed procedures of Palette can be divided into the following three steps. First, spot clusters are defined in the ST data and the proportion of each defined cluster is inferred in bulk RNA-seq data (Fig. 1a). Specifically, highly expressed genes in both ST and bulk RNA-seq data are used for spatial clustering of the ST data. Here, BayesSpace^20^, and MuSiC^16^ are employed for deconvolution to estimate cluster abundances in bulk data. This step can effectively eliminate the batch effect caused by technical differences in sampling, mRNA capture, platform, etc. between the two experiments. Second, a variable factor is introduced to adjust the cluster expression matrix (Fig. 1b). To obtain the variation of each gene in ST and bulk data, a pseudo bulk vector is achieved as the cross product of the cluster expression matrix of ST data and the cluster proportions of bulk data, so that the variable factor vector can be calculated by the ratio of the input bulk to the pseudo bulk vector. Consequently, the stable genes and variable genes can be distinguished by the distribution of the variable factor vector. The adjusted matrix is obtained by taking the dot product of the cluster expression matrix of ST slice and the variable factor. This step can effectively overcome the common sparsity problem in spatial transcriptomics technologies, and the adjusted matrix not only contains the cluster composition information but also fully retains the accuracy of the bulk transcriptome in the detection of lowly expressed genes. Third, the expression of each spot is estimated through an iteration algorithm (Fig. 1c). In each iteration, the procedure begins by selecting one random spot (i) and its nearest neighbouring spots (Local i). The expression of spots belonging to the same cluster is aggregated to form a pseudo-cluster expression data called local ST (LST). Assuming the ratio of LST to the reference ST data is equal to the ratio of the adjusted LST to the adjusted matrix derived from the previous step, the expression of adjusted LST can be calculated and evenly allocated into the selected spots of this cluster. The loop then proceeds to the next iteration (iteration j), and after multiple iterations, typically thousands of times, the average expression of each spot is almost stable, which is considered as the output estimated expression.

The expression patterns on Palette reconstructed ST show enhanced spatial specificity and continuity (Fig. 1c). Our algorithm incorporates spot characteristics and spot-spot distances, emphasizing cluster-specific expression, while leveraging expression from bulk data to adjust gene expression in the ST spots. Additionally, the assumption that the neighbouring spots of the same cluster share similar gene expression enables imputation based on gene expression in neighbouring spots. This strategy partially mitigates the limitation of low detected gene numbers in each spot. Overall, the Palette pipeline serves as a valuable tool for inferring spatial gene expression patterns from bulk RNA-seq data, striving to generate accurate predictions of spatial gene expression that closely resemble the expression patterns in bona fide tissues.

### Palette enables the prediction of gene expression patterns with higher spatial specificity and accuracy

To assessed the performance of Palette, we first utilized two consecutive slices (referred to slices 4 and 5) from the Stereo-seq data^13^ of *Drosophila* E14-16 (14-16 hours post egg laying) serial sections (Fig. 2a). We converted slice 5 into a pseudo bulk and used it as Palette’s input, with slice 4 serving as the ST reference. We observed that Palette-implemented slice 4 did not result in considerable changes in the molecule numbers of each spot, as the gene expression levels of slice 4 and slice 5 were similar. However, there was a significant increase in the feature numbers (gene numbers) of each spot (Fig. 2b). This increase was attributed to the supplementation by Palette, which leveraged the gene expression of neighbouring spots belonging to the same cluster.

**Fig. 2.**
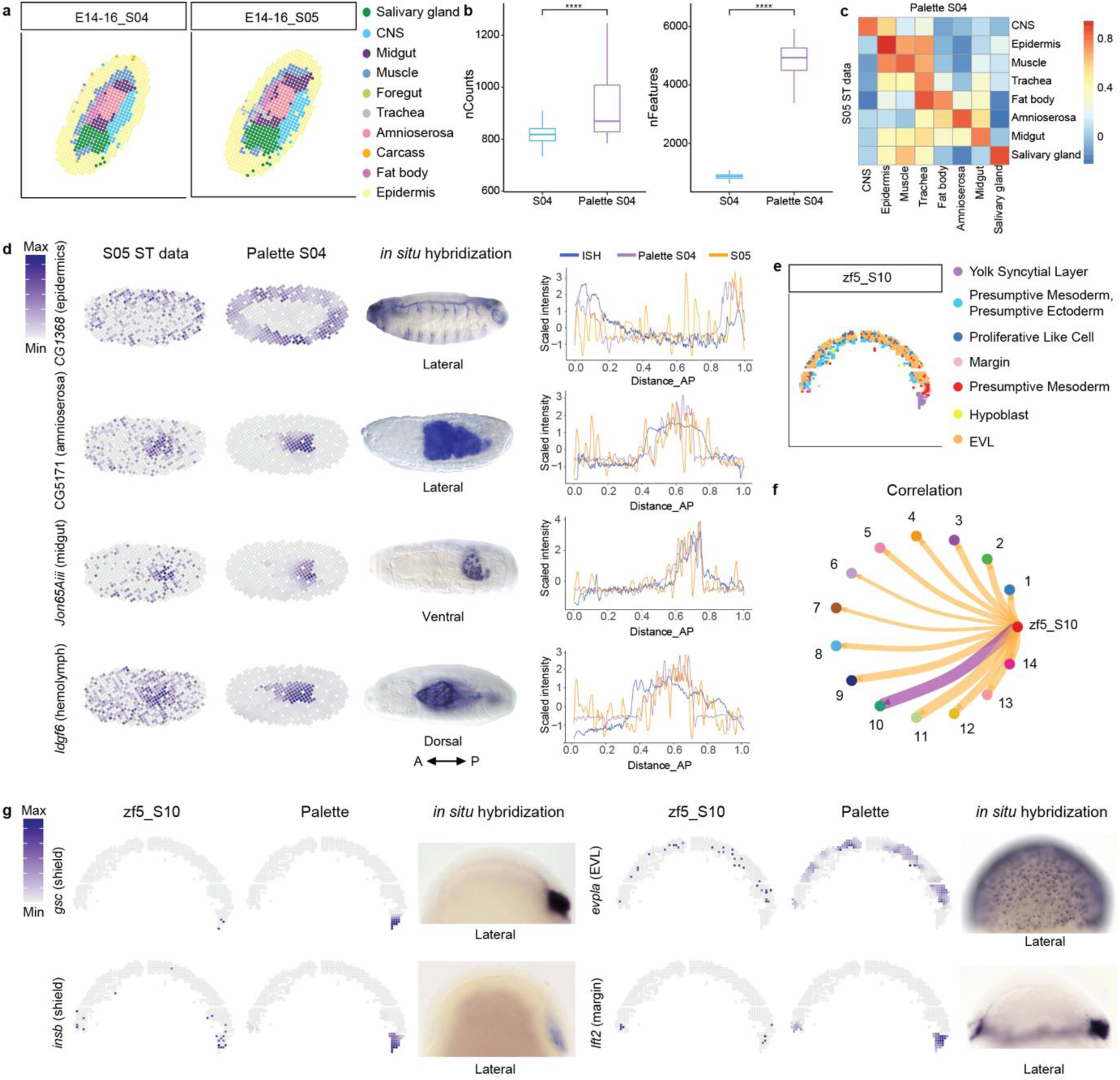
The implementation of Palette resulting in more specific gene expression patterns. **a,** The clustering and annotation of two adjacent slices from the Stereo-seq data^13^ of E14-16 *Drosophila* embryo. **b,** Boxplots showing the numbers of molecules and genes in each spot before and after implementing Palette. The substantial increase in gene number is due to the supplementation from neighbouring spots, based on the assumption that neighbouring spots within the same cluster exhibit similar gene expression patterns. **c,** Heatmap showing the expression correlation of marker genes for each cluster before and after implementing Palette. The colour bar represents the Pearson correlation coefficient with positive correlation in red and negative correlation in blue. **d,** Spatial expression patterns of marker genes on the *Drosophila* Stereo-seq slices. Intensity of colour represents the expression levels of each marker gene. For each gene, the spatial patterns from the Stereo-seq S05 slice and the Palette-implemented S04 slice are shown on the left, and the ISH images from BDGP database are shown in the middle. The intensities of signals along the AP axis are shown on the right. **e,** The clustering and annotation of the selected slice from the Stereo-seq data^12^ of 5.25 hpf zebrafish embryo. **f,** Circle plot showing the expression correlation network between the serial bulk data of 6 hpf zebrafish embryo and the pseudo bulk of the Stereo-seq slice. Stroke weight indicates the strength of the Pearson correlation coefficient. **g,** Palette inferring spatial expression patterns of 6 hpf zebrafish embryo bulk data on the 5.25 hpf zebrafish Stereo-seq slice. Since zebrafish embryos at 5.25 hpf and 6 hpf exhibited similar expression patterns, we used Palette to infer spatial gene expression from the 6 hpf zebrafish embryo bulk data using the 5.25 hpf ST data as a reference. Intensity of colour represents the gene expression levels. For each gene, the spatial patterns from the Stereo-seq S10 slice and the Palette-implemented S10 slice are shown on the left, and the correlated ISH images shown on the right are from ZFIN and published data^21, 22^.

Furthermore, we observed strong correlations in the expression of top marker genes between the same annotated clusters of Palette-implemented slice 4 and slice 5 ST data (Fig. 2c), indicating that Palette successfully preserved the molecular characteristics of each spot. Notably, Palette-implemented slice 4 exhibited similar gene expression patterns to the slice 5 ST data, with these patterns being even more spatially specific and closely resembling the *in vivo* expression patterns observed through ISH (Fig. 2d, Fig. S1a and Fig. S1b). These results suggest that Palette’s ability of gene supplementation contributed to improved continuity in the expression patterns of the implemented slices. Moreover, since Palette considered the gene expression levels within each cluster, genes with highly differential expression among clusters exhibited more specific expression patterns.

To further evaluate Palette’s performance, we applied it to two additional datasets of zebrafish embryos^3, 12^: we selected one middle slice from a Stereo-seq data as the ST reference (Fig. 2e), and the slice 10 from a bulk data was selected as the corresponding input slice based on a correlation test (Fig. 2f). We then compared the expression pattern of genes on the original ST slice and the Palette-implemented slice (Fig. 2g and Fig. S1c). It was evident that the Palette-implemented slice exhibited more spatially specific expression patterns, which were more similar to the patterns observed through ISH.

Overall, Palette successfully inferred spatial gene expression from the bulk data of real biological samples, generating expression patterns with improved continuity and higher spatial specificity.

### Using Palette to infer spatial gene expression from bulk RNA-seq data of zebrafish serial cryosections

To generate a more precise 3D ST dataset of zebrafish embryos, we first performed serial cryosections of embryos at three developmental stages along the left-right axis and conducted high-depth bulk RNA-seq (Fig. 3a, Fig. 3b and Fig. S2). Then, Palette was applied to create a more accurate zebrafish spatial transcriptomic atlas.

**Fig. 3.**
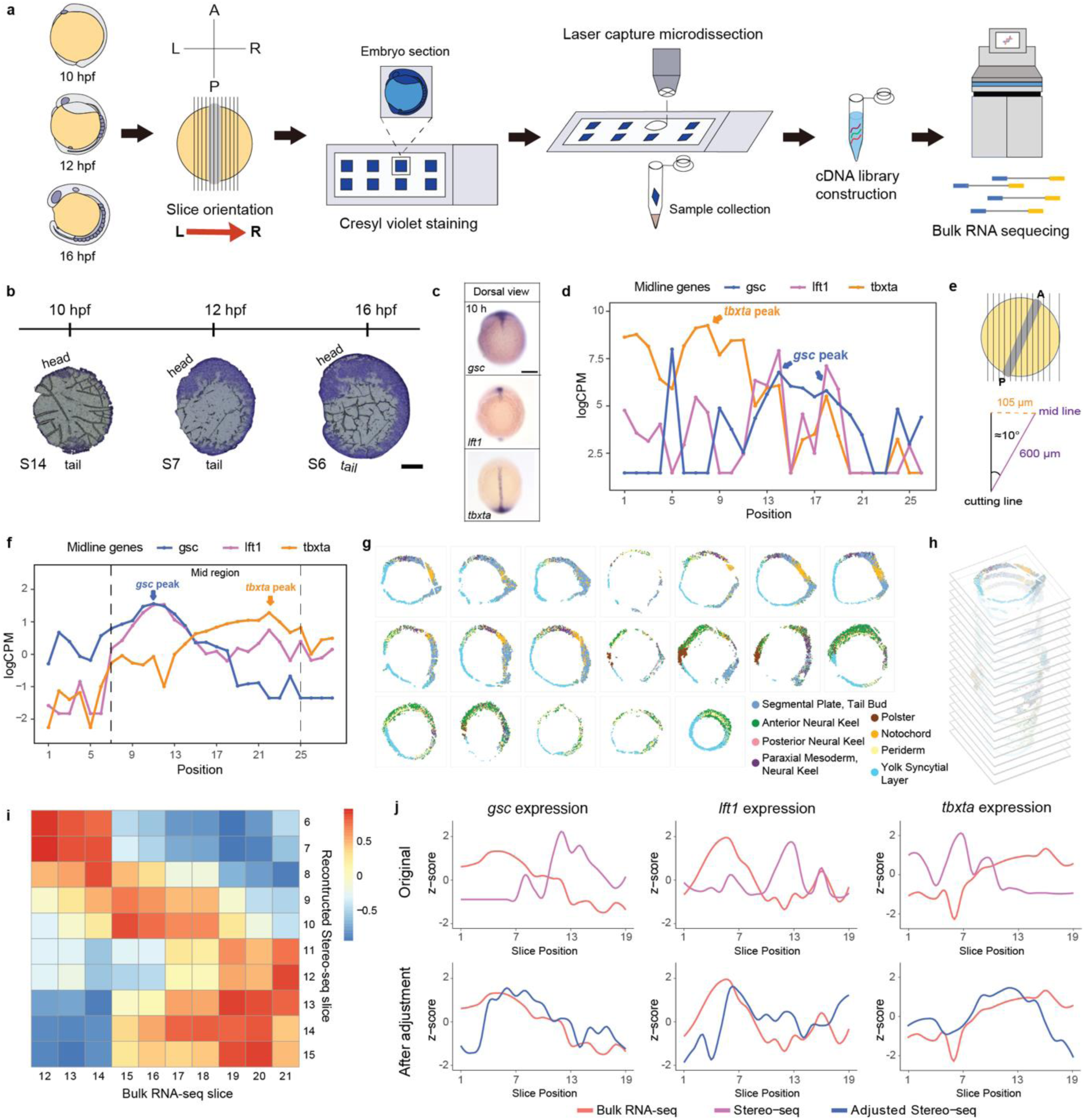
Processing the serial bulk RNA-seq data and the Stereo-seq data. **a**, Schematic representation of the workflow for generating the serial bulk RNA-seq data of zebrafish embryos. **b**, Cresyl violet staining of the cryosectioned slices. Each slice is 20 µm thick. The regions stained by cresyl violet correspond to cells. **c**, ISH images showing the expression patterns of midline genes, *gsc*, *lft1* and *tbxta*, from dorsal view. **d**, **f**, Expression plot showing the expression patterns of midline genes along left-right axis in the Stereo-seq data (d) and the bulk RNA-seq data (f). **e**, Diagram illustrating the midline of the Stereo-seq data tilted towards the left. **g**, The adjusted Stereo-seq slices. Poor-quality and severely damaged slices were discarded. Each spot is labelled with cell type annotations. **h**, 3D construction of the Stereo-seq slices. **i**, Correlation heatmap between the bulk RNA-seq data and the re-segmented Stereo-seq slices. The colour bar represents the Pearson correlation coefficient. **j**, Expression plots showing the comparison of gene expression patterns before and after slice alignment. Scale bar: 100 µm (b), 200 µm (c).

Before implementing Palette, we aligned the ST data with the bulk RNA-seq data using three midline genes—*gsc*, *lft1*, and *tbxta*—as metrics of alignment accuracy (Fig. 3c). Analysis revealed that the slice cutting lines were not parallel to the embryonic midline in both the Stereo-seq and our bulk RNA-seq data (Fig. 3d and Fig. 3f). The anterior midline gene *gsc* and the posterior midline gene *tbxta* appeared on different slices, and the tilt directions differed between the Stereo-seq data and the bulk RNA-seq data, as indicated by the positional relationships of *gsc* and *tbxta* (Fig. 3d-f) along the left-to-right direction.

To align these two datasets, we first adjusted and orientated the ST slices (Fig. 3g). We then overlaid them sequentially at consistent intervals (Fig. 3h), creating a 3D ST dataset that could be rotated and re-segmented to facilitate alignment. The efficacy of alignment was evaluated using a correlation coefficient derived from the expression patterns of genes with known AP differentiation (See Methods). Through continuous adjustments—rotating, re-segmenting, and recalculating correlations—we identified the configuration with the highest mean correlation coefficient. This configuration was deemed optimal for aligning the re-segmented slices with those from the bulk RNA-seq (Fig. 3i). The expression patterns of midline genes in the re-segmented Stereo-seq slices closely aligned with those in the bulk RNA-seq slices (Fig. 3j).

### Integrating zebrafish spatial transcriptomics data and imaging data to construct *Dre*STEP

Palette was applied to reconstruct a 3D zebrafish ST atlas. However, the ST sections exhibited extrusion and deformation (Fig. 3g and Fig. 3h), resulting in spatial distortions. To generate a 3D ST atlas that enables accurate visualization of gene expression patterns within zebrafish embryos while preserving their comprehensive morphology, we projected the ST spots onto 3D zebrafish embryo imaging data^19^. This approach utilized the detailed morphological representation provided by the 3D imaging, where each cell is assigned a spatial coordinate, serving as a precise reference for the projection of ST spots.

Prior to spot projection, the ST data and the 3D imaging data was initially aligned. We scaled the embryo to similar sizes in both datasets, and selected three spots located at the head, tail and middle of the midline from each dataset. These three paired spots were then utilized for alignment using the Kabsch algorithm^23, 24^, which is a method for calculating the optimal rotation matrix that minimizes the root mean squared deviation (RMSD) between two paired sets of spots. This resulted in the alignment between the ST data and the imaging data (Fig. 4a).

**Fig. 4.**
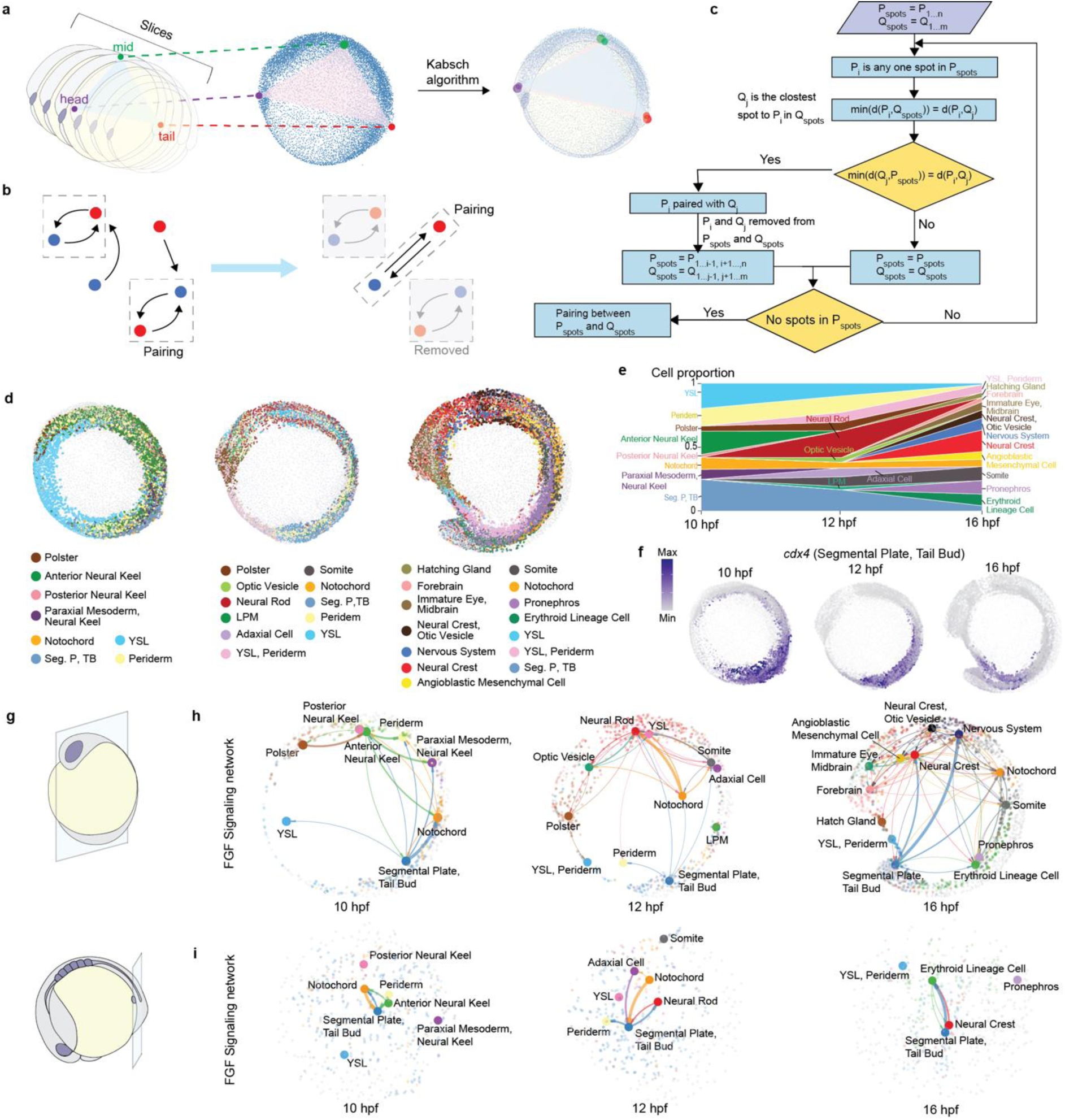
Projection of ST spots on 3D images and analysis of spatial cell-cell communication. **a,** Diagram showing the overall alignment between the ST data and the imaging data using the Kabsch algorithm. **b,** Diagram indicating the pairing principle for the ST coordinates and the imaging coordinates. In each interaction, the ST spot and the imaging spot closest to each other are paired, which is considered as the optimal solution of this interaction. Paired spots are removed from subsequent interactions, and the loop continues until each ST spot is paired with an imaging spot. **c,** Flow chart showing the process of the pairing. **d,** Lateral view of *Dre*STEP. Each spot is coloured with cell type annotation. **e,** Stacked area plot showing cell proportions in each stage. **f,** Expression patterns of *cdx4* on *Dre*STEP. Intensity of colour represents the gene expression levels. **g,** Schematic showing the *in silico* sections cut for spatial cell-cell communication analysis. **h, i,** Analysis of FGF signalling pathway network in the midline (h) and tail sections (i). Each spot is coloured with cell type annotation. The stroke weights indicate the interaction strength. YSL: Yolk syncytial layer; LPM: Lateral plate mesoderm; Seg P, TB: Segmental plate, Tail bud.

Following the alignment, the projection from ST spots to imaging spots was achieved using a loop algorithm inspired by the conception of Greedy algorithm^25^ (Fig. 4b and Fig. 4c). The entire process resulted in the spatial gene expression atlas of zebrafish embryos of three developmental stages, which was named *Dre*STEP (Fig. 4d). *Dre*STEP encompassed zebrafish embryos at 10 hpf, 12 hpf and 16 hpf, and these stages corresponded to post-gastrulation and tail elongation processes. Consequently, *Dre*STEP precisely allocated cell clusters and gene expression on a bona fide zebrafish embryo with 3D coordinates (Figs. 4d-4f).

### Exploration of spatial cell-cell interactions in *Dre*STEP

*Dre*STEP enables the visualization of gene expression patterns in 3D view of zebrafish embryos, along with their comprehensive morphology (Fig. 4f and Figs. S4a-S4c), which allows for the freewheeling selection of specific regions of embryos for spatial cell-cell interaction (CCI) analysis.

We extracted the midline and tail sections from *Dre*STEP and employed CellChat^26, 27^ for spatial cell-cell communication analysis (Fig. 4g). At 10 hpf, we observed strong interactions between tail bud/segmental plate cells and notochord cells in both sections, with the FGF signalling pathway playing a significant role in mediating this interaction (Fig. 4h and Fig. 4i). Additionally, tail bud/segmental plate cells were found to send FGF signals to neural cells, and these cell-cell interactions persisted at 12 hpf and 16 hpf, with tail bud/segmental plate cells continuing to send FGF signals to both notochord and neural cells (Fig. 4h and Fig. 4i). Notably, the strength of FGF signalling from tail bud/segmental plate cells to neural cells increased at 16 hpf. These results indicated that tail bud/segmental plate cells served as a strong FGF signalling centre regulating neighbouring cells, which can be evidenced by the reported roles of FGF signalling in somite development^28–30^, caudal spinal cord development^31^ and posterior notochord development^30^.

Beyond FGF signalling, we also observed that other signalling pathways significantly contributed to the cell-cell interactions in the midline and tail sections. Throughout all three developmental stages, we detected strong interactions between neural cells with neighbouring cells through Wnt/β-catenin signalling (Fig. S4d). Additionally, we found that tail bud/segmental plate cells consistently emitted BMP signals to adjacent cells including notochord, adaxial, and erythroid lineage cells (Fig. S4e). These findings aligned with prior knowledge indicating that Wnt/β-catenin signalling was involved in regulating the neural plate patterning^32^, and BMP signalling was activated in tail bud region, contributing to tail formation^33^.

Our spatial cell-cell communication analysis suggests that different morphogens mediated diverse CCIs during embryonic development. The complex cellular networks formed by these CCIs may guide the formation of organ collectives and ensure the robust of organogenesis. In summary, *Dre*STEP proves to be an excellent zebrafish spatial atlas for visualizing gene expression patterns and investigating CCIs in specific regions of the embryo.

### Investigating morphogen distributions and cell fate specification in *Dre*STEP

During embryonic development, a group of signalling molecules, known as morphogens diffuse from localized sources, forming concentration gradients that provide spatial information to responding cells and guide their differentiation^34, 35^. The intersections of different morphogens with antiparallel gradients generate diverse cell types, contributing to the formation of precise patterns and structures^36, 37^ (Fig. 5a).

**Fig. 5.**
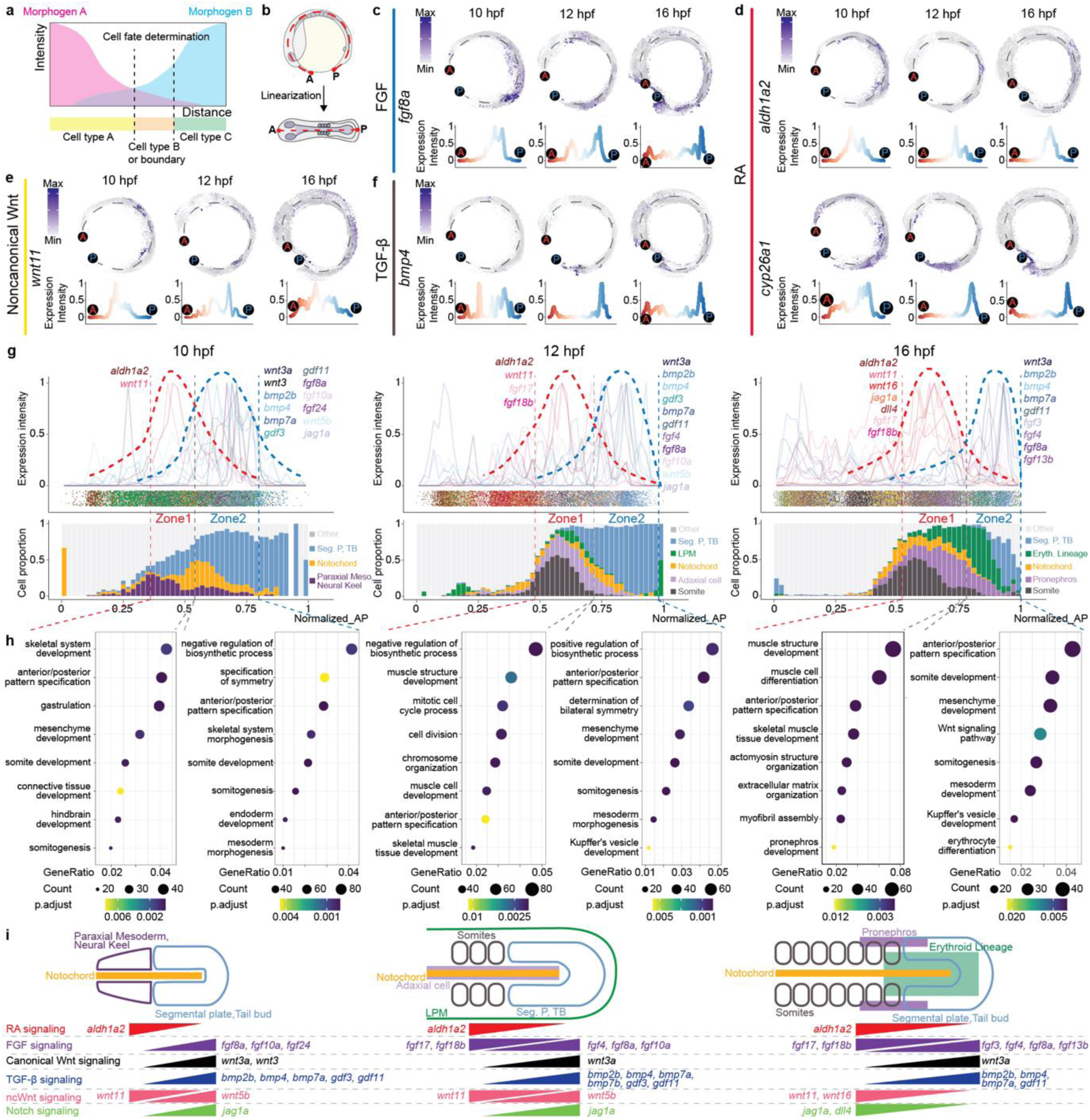
Morphogen gradients regulate the establishment of AP axis. **a,** Schematic diagram showing that role of antiparallel morphogen gradients in governing cell fate determination. **b,** Schematic diagram showing the linearization of *Dre*STEP. **c-f**, Plots displaying the expression patterns and intensities of representative ligands along the AP axis in FGF (c), RA (d), noncanonical Wnt (e) and TGF-β (f) signalling. The selected ligands show differential expression patterns in Zone1 and Zone2. **g,** Plots of gene expression intensities, cell types and cell type proportions along AP axis. The thick dashed lines in red and blue indicate the expression trends of trunk-enriched and tail-enriched genes; the thin dashed lines sperate Zone1 and Zone2 for GO enrichment analysis. **h,** Enriched GO terms in Zone1 and Zone2 respectively. **i,** Model diagrams showing the relationships between morphogen gradients and cell type specification in Zone1 and Zone2 at different developmental stages. Paraxial Meso: Paraxial mesoderm; LPM: Lateral plate mesoderm; Seg P, TB: Segmental plate, Tail bud; Eryth. Lineage: Erythroid Lineage; ncWnt: noncanonical Wnt.

The establishment of the AP axis involves the intricate interactions among morphogen gradients^22, 38, 39^. During tail elongation, morphogen gradients collectively regulate the extension and confinement of the AP axis, resulting in the precise specification and arrangement of tubular organ primordium along the body axis^38, 40, 41^. *Dre*STEP provides an appropriate platform to comprehensively analyse the expressing patterns of morphogens along the AP axis and investigate the relationships between the morphogen gradients and cell type distributions. We linearized the *Dre*STEP (Fig. 5b, See Methods) and focused on ligands involved in canonical Wnt, noncanonical Wnt, Notch, Sonic hedgehog (SHH), RA, FGF, and TGF-β signalling, visualizing their expression intensities along the linearized AP axis (Figs. 5c-f and Figs. S5-S8). We observed two adjacent regions along the linearized AP axis at all three time points, and each enriched with distinct group of ligands (Fig. 5g). These regions, designated as Zone1 and Zone2, were subjected to Gene Ontology (GO) enrichment analysis using sets of differentially expressed (DE) genes to investigate the functional characteristics of cells within each zone (Fig. 5g and Fig. 5h, Data S1-S9).

At the end of the gastrulation (10 hpf), tail elongation commenced with various cell types, beginning to be specified along the AP axis (Fig. 5g left). Notably, Zone1 consisted of paraxial mesoderm cells, while Zone2 predominantly comprised segmental plate/tail bud cells. GO enrichment analysis revealed terms related to somite development for both zones, such as “skeletal system”, “somite development”, and “somitogenesis” (Fig. 5h left). Furthermore, Zone2 encompassed the entire tail region, displaying GO terms associated with posterior development, such as “endoderm development” and “mesoderm morphogenesis”.

At 12 hpf and 16 hpf, as tail elongation progressed, more cell types were specified along the AP axis. The boundary between Zone1 and Zone2 shifted posteriorly. Zone1 primarily consisted of trunk region cells, such as somite cells, with GO terms related to muscle development, such as “muscle structure development”, “muscle cell development”, and “skeletal muscle tissue development”. Zone2 continued to predominantly consist of segmental plate/tail bud cells, with GO terms including “somitogenesis”, “somite development”, and “mesenchyme development”, indicating these cells’ high mobility and contribution to somitogenesis and tail elongation (Fig. 5h mid and right). The boundary between Zone1 and Zone2 coincided with the position of somite cells, highlighting the essential roles of antiparallel morphogen gradients in somitogenesis. Additionally, pronephros and erythroid lineage cells were specified at 16 hpf and distributed in both Zone1 and Zone2, with Zone1 containing a higher proportion of pronephros cells and Zone2 exhibiting a higher abundance of erythroid lineage cells (Fig. 5g).

Based on the observed transcriptional morphogen gradients and cell type distributions along AP axis in Zone1 and Zone2 at the three developmental stages, we created a diagram to summarize those findings (Fig. 5i). Assuming that the transcriptional level of a morphogen reflects its activity level, our model demonstrated the presentence of opposing concentration gradients, which could guide the cell type specification along the AP axis. Our analysis showed that Zone1 enriched aldehyde dehydrogenase *aldh1a2* (Fig. 5d); while Zone2 showed a high expression of *wnt3a* and *fgf8a* (Fig. 5c and Fig. S5). These observations were consistent with previous studies^42–46^ demonstrating the role of anterior RA signalling and posterior Wnt&FGF signalling in establishing the determination front of newly formed somites. In addition to these ligands, Zone2 exhibited enrichment of other FGF ligands, such as *fgf10a*, *fgf4* and *fgf13b* (Fig. 5g), suggesting their collective roles in regulating zebrafish embryonic posterior development. Interestingly, Zone1 also showed enrichment of certain FGF ligands, including *fgf17* and *fgf18b* (Fig. 5g), suggesting another FGF signalling cascade probably participated in somite development. Moreover, Zone2 enriched the ligands associated with TGF-β signalling, including *bmp2b*, *bmp4*, *bmp7a* and *gdf11*, aligned with the well-studied roles of BMP signalling in tail development^47–50^. Zone1 and Zone2 also exhibited enrichment of different noncanonical Wnt signalling ligands, *wnt11* and *wnt5b*, respectively, suggesting their important roles in regulating the pattern formation in these zones. Another interesting observation was that the expression of the Notch signalling ligand *jag1a* shifted from high expression in Zone2 at 10 hpf and 12 hpf to high expression in Zone1, along with *dll4*, at 16 hpf, suggesting changes in the zones where Notch signalling functions during zebrafish development.

In summary, our work systematically assessed the dynamic transcriptional profiles of morphogens along the AP axis and highlighted the interactions between adjacent zones exhibiting antiparallel morphogen gradients. These findings underscored the crucial roles of these morphogens in orchestrating pattern formation during zebrafish development, laying the foundation for investigating the regulation of AP refinement in further studies.

### Identification of key transcriptional regulatory cascades during the AP axis canalization

Diverse morphogen signals intricately interact to instruct the refinement of AP axis, involving cross-regulation of their intracellular pathways and downstream transcription factors (TFs)^51, 52^. To identify key morphogens and their downstream TFs that are essential for accurately determining the AP fate of various cell types within the embryo, we employed a random forest model (Fig. 6a), where we considered the expression levels of all morphogens and TFs as variable factors, to identify which ones are crucial for establishing the AP identities. We found that several morphogen ligands from the FGF, Wnt, RA, Notch and BMP signalling pathways as key determinants of AP identities (Fig. 6b and Fig. S9). The regulatory potential of these morphogens in AP axis formation have been substantiated by previous studies^47–50, 53, 54^, reinforcing the validity of our approach.

**Fig. 6.**
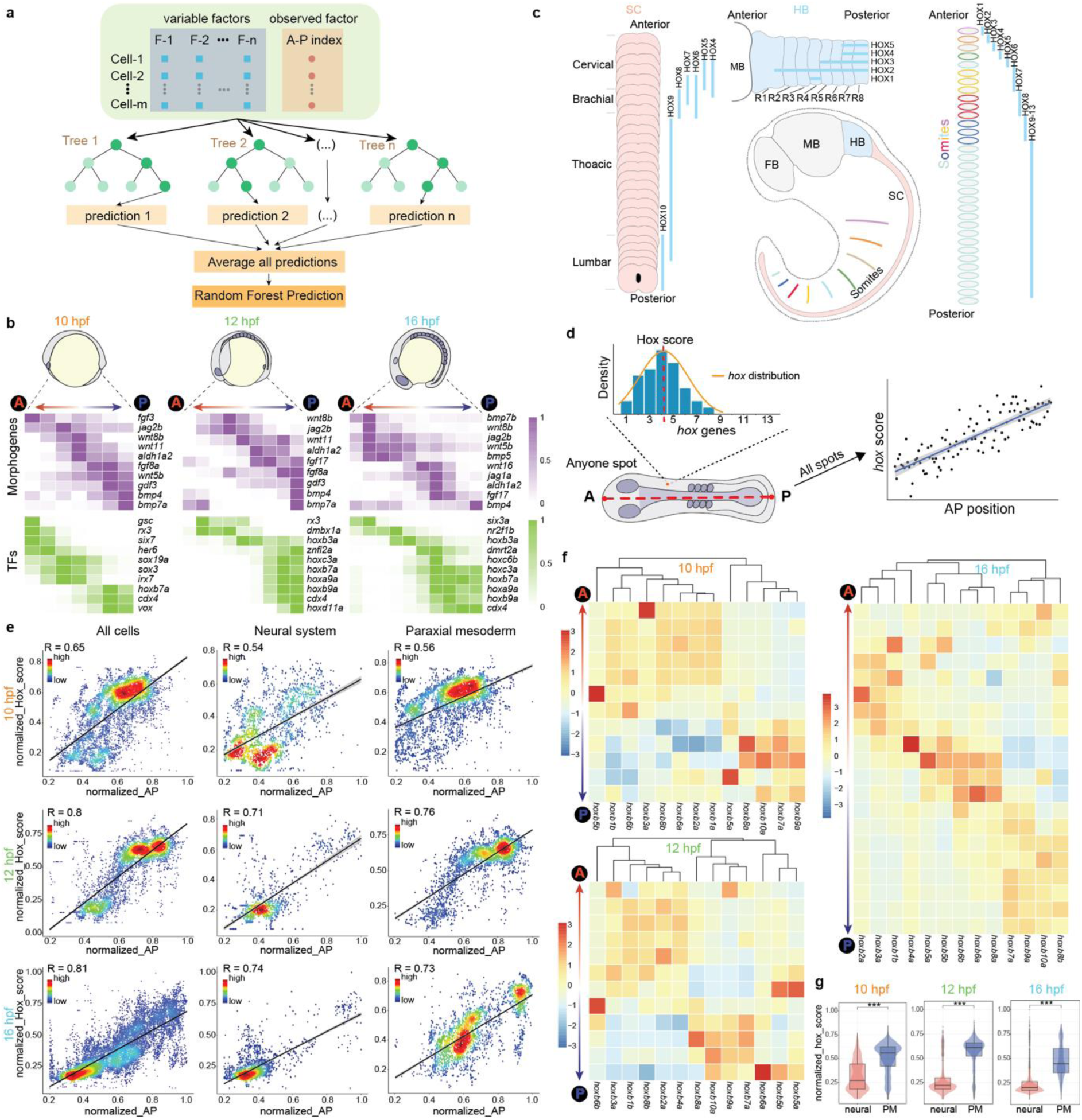
Establishment of AP axis. **a,** Schematic diagram showing the training for the random forest model. **b,** Heatmaps showing the expression intensities of key morphogens and transcription factors along AP axis at 10 hpf, 12 hpf and 16 hpf. **c,** Schematic diagram illustrating the spatial distribution of *hox* genes in neural and somatic systems. **d,** Schematic diagram showing the calculation of the Hox score of each cell and the assessment of correlations between Hox scores and AP positions. **e,** The correlations between Hox scores and physical AP positions. Colour indicates the spot density, with high in red and low in blue. **f,** Heatmaps showing the expression levels of *hoxb* genes along AP axis at 10 hpf, 12 hpf and 16 hpf. Intensity of colour represents z-score with high in red and low in blue. **g,** The Hox score distributions in neural system and paraxial mesoderm. Colour scale: Expression intensity (b), Z-score (f).

Interestingly, ligands within the same signalling pathway can exhibit distinct distributions along AP axis, suggesting their dominant roles in refining AP axis at different developmental stages. For instance, *fgf3*, anteriorly distributed, was a key morphogen at the 10 hpf stage, while *fgf17* and *fgf8a*, located in the posterior trunk and tail regions, respectively, were critical at both the 12 hpf and 16 hpf stages (Fig. 6b). This emphasizes the need to interpret morphogen gradients within a spatiotemporal context.

Among the identified key TFs, the Hox gene family genes emerged as significant regulators, underscoring their vital roles in AP axis regulation (Fig. 6b and Fig. S10). The Hox genes, a subset of conserved homeobox genes, exhibit both temporal collinearity and spatial collinearity in their expression, allowing them to specify regions along the AP axis and contribute to body plan formation^55–57^ (Fig. 6c). To further investigate the relationships between Hox gene expression and AP axis refinement, we defined a “Hox score”, which serves as an estimation for the most probable Hox gene expressed in each spot (Fig. 6d). We performed correlation analysis between Hox scores and physical AP identities across three developmental stages (Fig. 6e). Our results revealed a positive correlation between the Hox score and physical AP identities, with this correlation strengthening as development progressed. This trend was consistently observed in both the neural system and paraxial mesoderm, where Hox genes were independently expressed along the AP axis.

Interestingly, the neural system exhibited a lower Hox score compared to paraxial mesoderm (Fig. 6g), suggesting a “time discrepancy” between these two systems in canalizing their AP “avenue”. These observations highlight the increasingly significant regulatory role of Hox genes in AP axis refinement. Furthermore, we examined the expression patterns of four Hox clusters: *hoxa*, *hoxb*, *hoxc*, and *hoxd*, along the AP axis (Fig. 6f and Fig. S11). The *hoxb* cluster exhibited the most pronounced correlation with the physical AP identities, suggesting that the *hoxb* family genes may serve as master regulators in refining the AP identities during development.

## Discussion

In this study, we present Palette, a pipeline that utilize existing ST data as the only reference to infer precise spatial gene expression patterns from bulk RNA-seq data. The gene expression patterns predicted by Palette exhibit enhanced spatial continuity and improved spatial specificity, closely resembling experimentally observed patterns (Figs. 2d, 2g and Figs. S1b, S1c). Furthermore, Palette can incorporate spot characteristics directly from histological images of tissue slices, enabling more accurate spot characterization compared to reliance on spatial clustering alone (Fig. S12b). Palette is also applicable to comparative analyses of spatial gene expression patterns in different conditions, as demonstrated using human pancreatic ductal adenocarcinoma (PDAC) data^58^. Here, Palette inferred spatial gene expression patterns from bulk RNA-seq datasets^59^ of normal and tumour tissue slices, revealing a notable decrease in tumour-specific gene expression in the normal tissue slice (Fig. S12). Ongoing research aims to expand Palette’s application scenarios and explore further possibilities for analysing and interpreting spatial gene expression patterns.

Leveraging the capabilities of Palette, we constructed a comprehensive spatial gene expression atlas, *Dre*STEP, by integrating transcriptomics from serial sections and 3D images of zebrafish embryos^19^. *Dre*STEP not only facilitates the visualization of gene expression patterns in 3D morphology of zebrafish embryos, but also allowed for the flexible selection of sections for spatial cell-cell interaction analyses. As a 3D spatial gene expression atlas, *Dre*STEP holds great potential for studying the intricate 3D spatial cell-cell interactions during zebrafish development.

We utilized a linearized version of *Dre*STEP to investigate the relationships between morphogen distributions and cell type specification along the AP axis during development. We identified two adjacent zones with antiparallel morphogen gradients, with the boundary of these two zones appearing to act as the determinant front for somite formation.

In addition, by employing a random forest model, we explored the correlations between morphogen/TF expression and the developing AP patterns. This analysis identified critical morphogens and downstream TFs essential for determining AP position at different developmental stages. Notably, the Hox family genes were identified as dominant TFs, with strong correlations between the expression patterns of *hox* genes and the cell physical AP positions. Importantly, our findings suggest that *hoxb* cluster likely plays a more significant role in AP axis formation compared to other *hox* clusters.

During the development of *Dre*STEP, we encountered several limitations that warrant future improvements. Firstly, the manual adjustment and alignment of Stereo-seq slices during 3D ST data construction were labour-intensive and introduce potential bias. Although tools like PASTE^60^ were employed, their performance was unsatisfactory, possibly due to the hollow circle shape of ST slices leading to tilted alignments. Newly developed tools such as STitch3D^61^ and Spateo^62^ could be worth exploring for assisting the alignment. Secondly, the Palette algorithm currently excluded genes not detected in any spot of the ST data. Leveraging serial bulk RNA-seq data, could enable the construction of gene co-expression networks along the cutting direction. This approach has the potential to assist in predicting the expression patterns in ST data. Thirdly, the performance of Palette and *Dre*STEP heavily relied on the quality of ST data. In this study, for example, the Stereo-seq data of 12 hpf zebrafish embryo had fewer slices on the right side (Fig. S3b), resulting in more blank spots in the right part of *Dre*STEP for the 12 hpf embryo. Therefore, with the development and improvements in ST techniques, Palette and *Dre*STEP will have even greater potential for analysing spatiotemporal gene expression.

## Methods

### Animal ethics

Wild-type zebrafish strain was maintained following the standard procedures, and the experimental procedures were approved by the Institutional Review Board of Zhejiang University. The approval number is ZJU20220375.

### The Palette algorithm

ST data was used as the reference for inferring spatial gene expression from bulk RNA-seq data. The input bulk expression matrix is **S** ∈ R*^n×1^*, which contains the expression information of *n* genes. BayesSpace^20^ was first employed to perform spatial clustering on the ST data using the genes highly expressed in both the ST data and the bulk data. There are *m* clusters identified through spatial clustering, and the average expression of each cluster is **C** ∈ R*^n×m^*. MuSiC^16^ was then employed to obtain the proportions of each defined cluster in the bulk data, **A** ∈ R*^m×1^*, through deconvolution. A pseudo bulk vector, **P** ∈ R*^n×1^* is constructed by taking the cross product of the cluster expression matrix of ST data and the cluster abundances of bulk data.

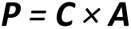

Each gene is assigned with a variable factor to adjust its expression. The variable factor vector, **K** ∈ R*^n×1^*, can be calculated as the ratio of the input bulk to the pseudo bulk vector.

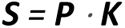

The adjusted matrix **M** ∈ R*^n×m^* is generated by the dot product of the cluster expression matrix of the ST slice and the variable factor.

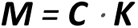

The pseudo bulk data of the reference ST slice is **T** ∈ R*^n×1^.* One random spot and its nearest neighbouring spots are selected, and the expression of spots belonging to the cluster *i* is aggregated to form a pseudo-cluster expression data LST, **L** ∈ R*^n×1^*. The regional cluster factor **R** ∈ R*^n×1^* is defined as the proportion of LST in the entire ST slice.

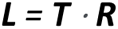

The ratio of LST to the reference ST data is equal to the ratio of the adjusted LST to the adjusted matrix, and thus the adjusted LST can be calculated by the dot product of the expression matrix of cluster *i* in the adjusted matrix **M^i^** and the regional cluster factor **R**. The expression of each spot in cluster *i*, **D** ∈ R*^n×1^*, is achieved through evenly allocation of the adjusted LST. **N** is the numbers of spots belonging to cluster *i* in this region.

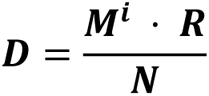

The average expression **D̅** ∈ R*^n×1^* after multiple iterations is considered as the estimated expression of the spot. Here ***p*** means that the spot has been selected for ***p*** times during iterations.

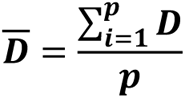

### Palette performance assessment on *Drosophila* slices

Two consecutive slices (referred to slices 4 and 5) were taken from the Stereo-seq data^13^ of *Drosophila* E14-16 serial sections. Given their adjacency, these two slices should exhibit similar gene expression levels and patterns. Slice 5 was then converted into a pseudo bulk and used as Palette’s input, with slice 4 serving as the ST reference. The predicted spatial expression patterns of slice 5 were compared to the actual ST data from slice and ISH images from BDGP database (www.fruitfly.org) for evaluating Palette’s performance.

### Palette performance assessment on zebrafish slices

One middle slice of zebrafish Stereo-seq data was selected as the ST reference^12^. A corresponded slice was selected as an input for Palette from the serial bulk RNA-seq data^3^ of zebrafish embryo using a correlation test. The correlation was calculated based on the expression of genes showing differential expression along the dorsal-ventral axis. The predicted expression patterns were compared to the ISH images from ZFIN (www.zfin.org) and published data^21, 22^ for evaluating Palette’s performance.

### Sample preparation for bulk RNA-seq

Live embryos at the required developmental stages of 10 hpf, 12 hpf and 16 hpf were rapidly embedded in optimal cutting temperature compound (OCT) and oriented in the bottom of steel embedding cassettes. Embedded embryos were rapidly frozen at −80°C for 10 minutes, and then transferred into a cryostat (Leica) at −20°C. In the cryostat, embedded embryos were removed from the steel embedding cassettes and cryosectioned at a thickness of 20 µm. Each slice was collected and placed on the PEN membrane slides (Leica) in the correct order. Membrane slides were stained using 1% (wt/vol) cresyl violet (dissolved in 70% ethanol) to roughly check cell number and ensure slice integrity. Each slice was extracted from membrane slides through laser capture microdissection system (Leica LMD6) and collected into a 1.5 mL Eppendorf tube containing 40 µL of PicoPure^TM^ lysis buffer (Thermo Fisher). Collected samples were incubated at 42°C for 30 minutes and then sent to Shanghai Ouyi Biology Medical Science Technology Co., Ltd. (Shanghai, China) for RNA extraction, cDNA library construction and sequencing. Paired-end sequencing at 150 bp read length was performed on a Novaseq 6000 instrument.

### Bulk RNA-seq data processing

Reads were aligned to the *Danio rerio* genome Ensembl Release 92 (GRCz11) using STAR v2.7.1a^63^. The aligned reads were assigned to each gene using featureCounts v1.6.0^64^. For each embryo, the gene counts of each slice were merged into a count matrix, and the genes that received more than 0.5 counts per million reads (CPM) in at least 3 slices were retained. The count matrices with slice position information were constructed into DGEList objects using edgeR^65–67^ for the following analysis.

### 3D embryo reconstruction

The spatiotemporal transcriptomics date used for 3D embryo reconstruction was obtained from the zebrafish Stereo-seq dataset^12^ available for download at https://db.cngb.org/stomics/zesta/download/. Each section was fitted into a 2D coordinate system, with the section centre serving as the origin. Before the reconstruction, severely broken slices and outlier spots were removed from the dataset. The position of each section on the 2D coordinate system was manually adjusted and aligned based on the section shapes and spot annotations. Additionally, the distance between neighbouring sections on the z-axis was estimated, and the corresponding z-axis coordinates were assigned to each section. By combining the spatial transcriptomics data and with the 3D coordinates, reconstructed embryos were generated.

### Alignment between bulk RNA-seq slices and re-segmented pseudo ST slices

Reconstructed embryos were rotated along x-axis and y-axis separately, and then re-segmented into section slices. Each of re-segmented slices was transformed into pseudo bulk data, which represented an aggregate expression profile for that slice. To determine embryo orientation, several genes exhibiting AP differential expression patterns were selected. The scaled expression levels of these genes across slices were generated for both bulk RNA-seq slices and pseudo bulk slices. To assess the alignment between bulk RNA-seq slices and pseudo bulk slices, Pearson correlation coefficients were calculated across these genes. The alignment with the highest mean Pearson correlation coefficient along the slices was considered as the optimal alignment.

### 3D projection of ST spots to imaging spots

The live imaging data of zebrafish embryos were obtained from the study by Shah et al., 2019^19^ available for download at https://idr.openmicroscopy.org under accession code idr0068. Both the centres of the ST data and the live imaging data were set as the origin, and the embryo size in the imaging data was scaled to a similar embryo size in the ST data. Three specific spots (head, mid of midline and tail) were selected from the both datasets. The Kabsch algorithm^23, 24^ was used to achieve the optimal alignment between the two paired sets using these three spot pairs. This algorithm involved rotating and transforming the coordinates of the three ST spots to align them with the corresponding spots in the live imaging data. By applying the rotation matrix obtained from the optimal alignment, the entire set of the ST spot coordinates was transformed. A transform matrix was obtained by calculating the difference between the coordinate of the head spot achieved from the spot alignment and the coordinate of the head spot after applying the rotation. The transform matrix was then applied to the coordinates of all the ST spots, which had undergone rotation. Through this process, the alignment between the ST data and the imaging data was achieved.

The pairing between the ST spots and the imaging spots was achieved through a looping algorithm based on the Greedy algorithm^25^. In each interaction of the loop, the ST spot and the imaging spot that were closest to each other were paired, which was considered as the optimal solution of the interaction. Paired spots were removed from subsequent interactions, while unpaired spots moved on to the next loop. The looping process continued until each ST spot was paired with an imaging spot. The expression information from the ST spots was then assigned to their corresponding imaging spots. The remained unpaired imaging spots were retained to preserve the overall morphology of the embryo.

### Spatial cell-cell communication analysis using CellChat

CellChat^26, 27^ was employed to analyse spatial cell-cell communication based on prior known zebrafish ligand-receptor interaction database CellChatDB. The section of interest was extracted from *Dre*STEP for analysis, which provided 2D section ST data. In the Stereo-seq data, each spot contained 15 × 15 DNA nanoball (DNB) spots. Consequently, in the section ST data, the spot diameter was set as 15, and the number of pixels spanning the spot size diameter was set as 225. The expression data of section ST data was pre-processed to identify over-expressed ligands and receptors for each cell group. Setting distance as constraints, CellChat inferred communication probability between two interacting cell groups. This inference was based on the average gene expression of a ligand in one cell group and the average gene expression of a receptor in another cell group. The communication probabilities of all ligands-receptors interactions associated with each pathway were summarized for analysis of the communication probabilities within in signalling pathways.

### Linearizing the AP axis in *Dre*STEP

The lateral view of *Dre*STEP was projected onto a 2D plane. The spots were fitted into a cycle. Each spot was then projected onto the cycle, with the projected spot representing the closest spot on the cycle to the original spot. By designating the most anterior spot as the origin, the AP position of each spot was determined by calculating the arc length from its projected spot to the origin. The AP value of each spot was then divided by the maximum AP value in the dataset to achieve the normalized AP value.

### Employing random forest model for prediction

For each cell, the normalized expression of morphogens or TFs was set as variable factors, while the cell’s normalized physical AP position was set as observed factor. We took 70% of the data to train a random forest model using the randomForest^68^ package. The importance of the variables to AP position was assessed by both increase in mean square error (IncMSE) and increase in node purity (IncNodePurity). Cross-validation was used to evaluate the number of variables. The top 6 important variables were selected, and the correlations between their expression and AP positions were visualized.

### Calculating Hox score of each spot

For each *hox* gene, its expression in each spot was divided by the maximum expression of that gene in the dataset, indicating the expression probability of that gene in that spot. Then, the expression of *hox* genes in spots can be converted to a repeated representation, where the number of repetitions corresponded to the expression probability of the gene in that spot. In our analysis, we made an assumption that the expression of *hox* genes in each spot followed a normal distribution. This assumption enabled generating a fitting curve of the normal distribution on the density plot of the *hox* genes. The Hox score was determined as the *hox* value at the peak of the normal distribution. The Hox score of each spot was then divided by thirteen, resulting in the normalized Hox score.

### *In situ* hybridization (ISH)

ISH was performed following the published protocol^69^. Embryos of required developmental stages were fixed in 4% PFA/PBS overnight at 4°C, and then transferred into 100% methanol (MeOH) for dehydration overnight at −20°C. Embryos were washed through 75%, 50% and 25% MeOH/PBST for 5 minutes each at room temperature and then three times for 5 minutes in PBST. Embryos older than 10 hpf were treated with proteinase K (10 μg/mL in PBST) for 30 s and then fixed in 4% PFA/PBS for 20 minutes at room temperature. Proteinase K treated embryos were washed four times for 5 minutes each in PBST. Embryos were transferred into Hybridization Mix (HM) and incubated at 70°C for 2-5 hours, and then the buffer was replaced by HM containing digoxigenin-11-UTP (Sigma-Aldrich, 11277073910) labelled probe. After overnight incubation at 70°C, embryos were washed through 75%, 50% and 25% HM/2xSSC at 70°C for 20 minutes each. Embryos were then washed in 2x SSC at 70°C for 20 minutes and washed in 0.2xSSC twice at 70°C for 40 minutes. 0.2xSSC were then progressively replaced by PBST at room temperature. Embryos were blocked in blocking buffer at 4°C for 3 hours, and incubated in blocking buffer with anti-DIG-AP antibody (1:10000 dilution, Sigma-Aldrich, 11093274910) at 4°C overnight on a low speed shaker. Embryos were washed 6 times for 15 minutes each in PBST to remove excess antibodies. Embryos were stained in NBT/BCIP staining solution, and the staining was stopped by washing twice in PBST when the expected staining patterns were observed.

## Data availability

The raw data of serial bulk RNA-seq has been deposited to the Gene Expression Omnibus (GEO) under accession number “<u>GSE262578</u>”. The published data used in this study can be accessed through the following links or accession number: (1) Stereo-seq data of *Drosophila* embryos^13^ (https://db.cngb.org/stomics/flysta3d/download/); (2) Stereo-seq data of zebrafish embryos^12^ (https://db.cngb.org/stomics/zesta/download/); (3) live imaging data of zebrafish embryos^19^ (idr0068 from https://idr.openmicroscopy.org); (4) Spatial transcriptomics data of human PDAC: GEO accession: “<u>GSE111672</u>”^58^; (5) Bulk RNA-seq data of human PDAC: GEO accession: “<u>GSE171485</u>”^59^.

## Code availability

The codes for Palette pipeline and bioinformatics analyses are deposited on GitHub (https://github.com/ldo2zju/DreSTEP). Any other custom code and data are available from the authors upon request.

## Additional information

Supplemental Information Supplementary Figures 1-12 and Supplementary Data 1-9.

## Acknowledgments

We thank Dr. Si-Yu He at Stanford University and the members of Laboratory of Development and Organogenesis (LDO) at Zhejiang University for helpful suggestions and discussions. This work was supported by grants from National Key Research and Development Program of China (2022YFA1103100), the National Natural Science Foundation of China (32050109, 32300677, 32300688, 82204772, U23A20513), the "Pioneer" and "Leading Goose" R&D Program of Zhejiang (2024C03106), and Ningbo Top Medical and Health Research Program (No. 2022030309).

## Author Contributions

PFX, YD, TC, XL, JL and XHF conceived and designed the research; YD, TC and JL designed Palette pipeline, constructed *Dre*STEP and performed analyses; XL, XXF, YH, XFY, LEY, HRL and ZWB performed experiments; YD and TC drafted the manuscript; JL, XHF and PFX edited the manuscript; all authors reviewed and approved the manuscript. PFX and XHF supervised this study. Fundings for the study were provided by PFX, YD, TC, JL and XHF.

